# A dynamical extracellular matrix coat regulates moruloid-blastuloid transitions of ovarian cancer spheroids

**DOI:** 10.1101/2020.07.03.186155

**Authors:** Jimpi Langthasa, Shruthi Narayanan, Rahul Bhagat, Annapurna Vadaparty, Ramray Bhat

## Abstract

Ovarian cancer metastasizes into the peritoneum through dissemination of transformed epithelia as multicellular spheroids ^1, 2^. Harvested from the malignant ascites of patients, spheroids exhibit startling features of organization typical to homeostatic glandular tissues^3^: lumen surrounded by smoothly contoured, adhered, and immotile epithelia. Herein, we demonstrate that cells of specific ovarian cancer lines in suspension, aggregate into dysmorphic solid ‘moruloid’ clusters that permit intercellular movement and penetration by new cells. Moruloid clusters can coalesce to form bigger clusters. Upon further culture, moruloid clusters mature into ‘blastuloid’ spheroids with smooth contours, lumen and immotile cells. Blastuloid spheroids neither coalesce nor allow penetration by new cells. Ultrastructural examination reveals a basement membrane-like matrix coat on the surface of blastuloid, but not moruloid, spheroids: immunocytochemistry confirms the presence of extracellular matrix proteins: Collagen IV and Laminin-322. Enzymatic debridement of the coat results in a reversible loss of lumen and contour. Debridement also allows spheroidal coalescence and cell intrusion in blastuloid spheroids and enhances adhesion to peritoneal substrata. Therefore, the dynamical matrix coat regulates both the morphogenesis of cancer spheroids and their adhesive interaction with their substrata, affecting ultimately the progression of the disease.

**Results:** Survival of women afflicted with epithelial ovarian cancer (EOC) trails behind other gynecological malignancies, despite improvements in surgical-pharmacological approaches^4,5^. The morbidity associated with the disease is a consequence of its transcoelomic route of metastasis: transformed epithelia of the fallopian tubes and ovaries in the form of spheroids, eventually home and adhere to the mesothelial lining of the peritoneum, occasionally invade through the underlying collagenous extracellular matrix and form secondary metastatic foci around abdominal organs1, ^6, 7^. EOC spheroids impede the drainage of the fluid from the peritoneal cavity and alter its composition; in turn the fluid, now known as malignant ascites serves as a pro-tumorigenic milieu for the spheroids^8, 9^

The formation and presence of spheroids within ascites of an ovarian cancer patient is strongly associated with recurrence of cancer and greater resistance to chemotherapy^10^. Therefore, in order to develop novel strategies to target spheroidal metastatic niche, it is essential to investigate mechanisms that underlie their morphogenesis. Several proteins have been proposed to mediate the adhesion between ovarian cancer epithelia that give rise to spheroids. These include transmembrane receptors such as CD44^11^, cell adhesion molecules, such as E-cadherin and N-cadherin^12^, matrix adhesion-inducing proteins such as integrins^13, 14^. Remarkably, a phase-contrast microscopic examination of spheroids from patients, or from aggregated epithelia of immortalized cancer lines cultured on low attachment substrata, shows features of morphogenetic organization: presence of a central lumen, radially arranged apposed epithelia and compacted surfaces. Such traits are cognate to organized morphogenesis within the glandular epithelial organs,^15^ which are built through principles that include, but are not limited to, cell-cell adhesion^16, 17^. In fact, loss of tissue architecture seen in tumorigenesis involves the disappearance of such morphogenetic traits (such as matrix adhesion and polarity)^18, 19^.

In this manuscript, we investigate how these traits are recapitulated in a fluid metastatic context. Using spheroids from patients with high grade serous adenocarcinoma and ovarian cancer cell lines, we show that the development of a basement membrane (BM)-like coat of extracellular matrix is responsible for the compaction and stability of cancer spheroids, for decreasing the motility of cells within it and for generation of lumen. The coat, which is rapidly replenished by cells upon enzymatic debridement, also prevents the attachment of spheroids to matrix substrata. This may have significant implications for the build-up of the massive cellular fraction within the malignant ascites of patients afflicted with ovarian cancer.

## Results

The cellular fraction obtained from the malignant ascites of patients with high grade serous ovarian cancers was isolated and cultured on low attachment substrata. When observed under phase contrast microscopy, we found both single cells and multicellular aggregates within the cellular fractions (Figure S1). The aggregates showed dysmorphic as well as in several cases, radially symmetric cellular architectures. Since such aggregates were mostly spherical in shape, they will henceforth be referred to as spheroids.

We asked whether the dysmorphic and spheroidal morphologies represent stages in a temporal spectrum of maturation of ovarian cancer MCAs, or do they represent interpatient or intrapatient lineages of cellular organization. Testing the former hypothesis required an assay, wherein the formation of clusters from cells could be temporally tracked and quantifiably assessed. We cultured four ovarian cancer lines: OVCAR3, SKOV3, OVCAR4 and G1M2 (an PDX-derived cell line from an Indian patient with high grade serous ovarian adenocarcinoma) on low attachment substrata. The morphologies of their aggregates (along with ascites-derived spheroids) was observed after 7 days. The spheroids were fixed and stained with DAPI and phalloidin to visualize their nuclei and F-actin cytoskeleton respectively using bright field- or confocal-microscopy (Figure 1A and B). Of the 4 cell lines, OVCAR3 and G1M2 consistently formed compacted spheroids with lumen, SKOV3 formed spheroids that did not show lumen formation (although smaller spheroids were observed with lumen) and OVCAR4 formed dysmorphic lumen-less multicellular clusters. Digitally sectioning OVCAR3 clusters along three orthogonal planes with laser confocal microscopy confirmed the lumen was indeed surrounded on all sides by cells (Figure S2). The surface of lumen containing aggregates examined by scanning electron microscopy (SEM) was smooth and compacted (Figure 1C). A cartoon depiction of the multicellular morphologies of spheroids from ascites and cell lines is shown in Figure 1D. Cell lines which gave rise to spheroids with lumen (Figure 1E) also showed a greater tendency to form more spherical aggregates (Figure 1F). A close examination of the inner lining of OVCAR3 and G1M2 spheroids revealed tightly apposed cancer epithelia forming a smooth lumen interior boundary, suggesting the lumen was a result of cellular organization that involved establishment of multicellular polarity (Figure 1G). To confirm this, we stained fixed spheroids for ZO-1, and found strong localization of the zonula occludens protein on both the outer and the luminal surface of spheroidal epithelia (Figure 1H), which is consistent with previous observations of inverted polarity in peritoneal spheroids of patients with colorectal carcinoma ^20^.

**Figure 1:**
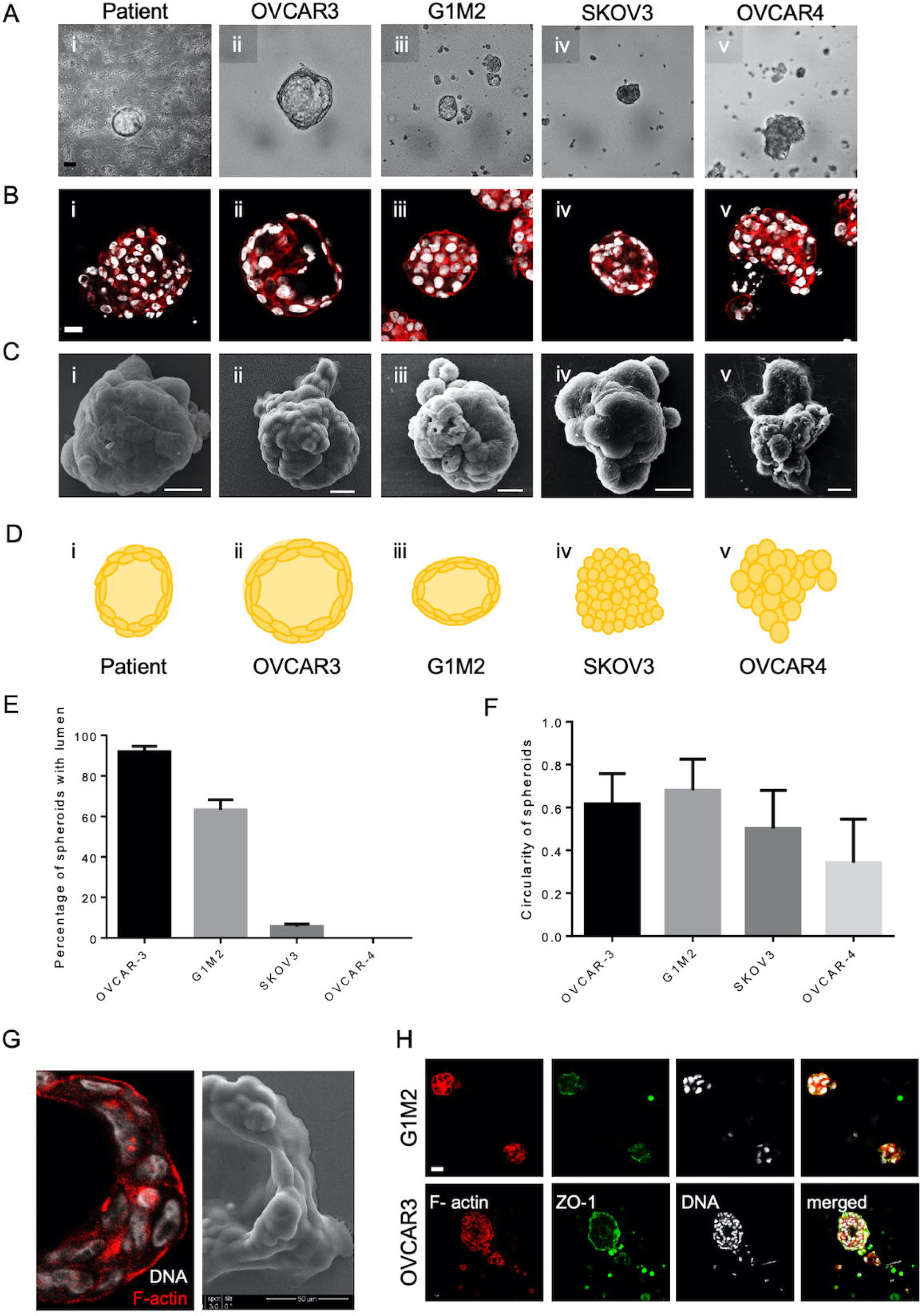
Spheroids of ovarian cancer epithelia show multicellular organization. (A-C) Photomicrographs of spheroids/clusters from patient malignant ascites (i), OVCAR3 (ii), G1M2 (iii), SKOV3 (iv) and OVCAR4 (v) cells imaged using phase contrast microscopy (A), laser confocal microscopy staining DNA (DAPI; white) and F-actin (Phalloidin; red)(B) and scanning electron microscopy (C). (D) Cartoon depiction of the morphology of spheroids/clusters, highlighting lumen formation and outer contour based on A-C. (E-F) Bar graphs showing the percentage of lumen-containing spheroids (E) and mean circularity of morphologies (F) derived from OVCAR3, G1M2, SKOV3 and OVCAR4 (G) Photomicrographs of OVCAR3 spheroids imaged using phase contrast (left) and scanning electron microscopy (right) showing tight apposition of cells that surround the spheroidal lumen. (J) Laser confocal photomicrographs of G1M2 (top) and OVCAR3 (bottom) spheroids stained for ZO-1 (green) F-actin (red) and DNA (white). Bars in E-F represent means+/-SEM. Scale bar for A, B and H is 50 μm and for Figure 1C is 10 μm.

Within cell lines wherein we observed lumen formation, the process of spheroid formation could be divided into an initial early step of cellular aggregation, followed by cavitation over a more protracted culture period. The morphology of early OVCAR3 and G1M2 spheroids (cultured for 1-2 days) assumed a more ‘moruloid’ appearance (no lumen, grape-like contour with cellular protuberances), whereas mature spheroids (cultured for 7 days) had a ‘blastuloid’ appearance with compacted surface and a lumen. The temporal distinction in these two morphologies was confirmed with bright field and phase contrast microscopy (Figure 2Ai-ii and 2Bi-ii), fluorescent labeling of DNA and F-actin (Figure 2Aiii-iv and 2Biii-iv), and scanning electron microscopy (Figure 2Av-vi and 2Bv-vi) in OVCAR3, G1M2, and to an extent in SKOV3 cells. SKOV3 cells formed smaller spheroids with lumen (Figure 2Ci-vi). The transition from moruloid to blastuloid phenotype was accompanied by an increase in size (Figure 2D and Figure S3i) and circularity (Figure 2E and Figure S3ii).

**Figure 2:**
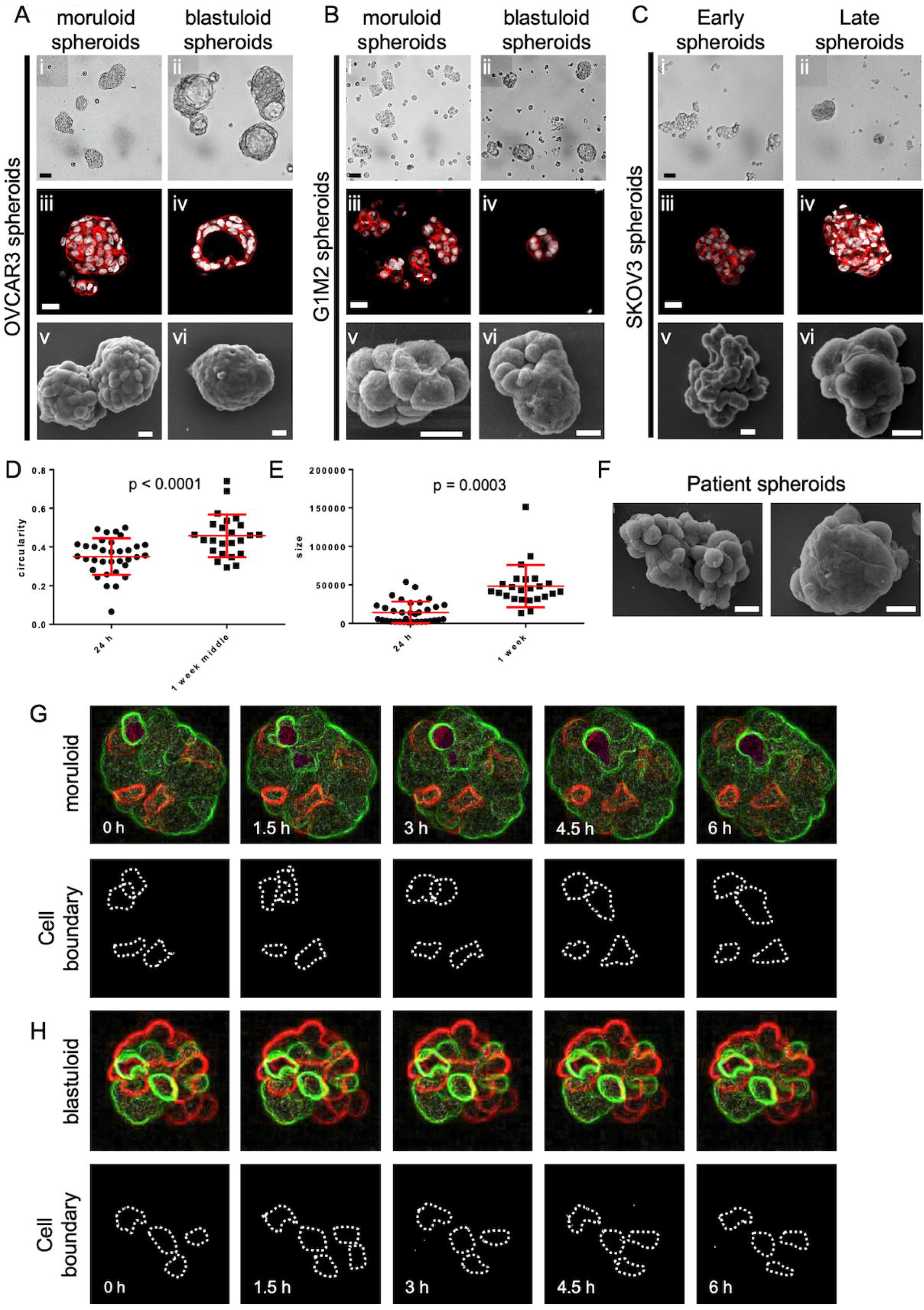
Distinctive features of moruloid and blastuloid morphologies in ovarian cancer spheroids. (A-C) Photomicrographs imaged using phase contrast- (i: moruloid, ii: blastuloid), laser confocal- (iii: moruloid, iv: blastuloid; staining for DNA using DAPI (white) and for F-actin using Phalloidin (red)), and scanning electron- microscopy (v: moruloid, vi: blastuloid) of OVCAR3 (A), G1M2 (B) and SKOV3 (C) spheroids (D-E) Graph showing change in size (D) and circularity (E) of OVCAR3 spheroids with moruloid (left) and blastuloid (right) morphologies. (F) SEM photomicrographs of patient-derived spheroids showing moruloid (left) and blastuloid (right) morphologies. (G) Photomicrographs taken at 0 hr, 1.5 hr, 3 hr, 4.5 hr and 6 hr from time-lapse laser confocal videography of moruloid spheroids (constituted from a suspension of GFP- and RFP-expressing OVCAR3 cells) showing rearrangement of motile cells within them (white dotted lines highlight the position of motile cells) (see Video S1) (H) Photomicrographs taken at 0 hr, 1.5 hr, 3 hr, 4.5 hr and 6 hr from time-lapse laser confocal videography of blastuloid spheroids (constituted from a suspension of GFP- and RFP-expressing OVCAR3 cells) showing rearrangement of motile cells within them (white dotted lines highlight the position of motile cells)(See Video S2) Error bars in D and E signify median with interquartile range. Significance was measured using unpaired t-test with Welch’s correction. Scale bar for A-C (i-iv) is 50 μm and for A-C (v-vi) and F is 10 μm Objective 20X.

Our proposition of a two-step morphogenesis of spheroids also found support when analyzing *ex vivo* ascites cell fractions. Both moruloid and blastuloid spheroids were found within the cellular fractions obtained from malignant ascites of patients (Figure 2F). The gradual conversion of a heterogeneous population to a more predominantly blastuloid one upon ex vivo culture suggested that a temporal progression of morphogenesis may also be taking place within disseminated spheroid niches *in vivo*. The presence of lumen within homeostatic epithelial architectures coincides with cellular immotility. We asked if the acquisition of blastuloid architecture is associated with a decrease in spheroidal cell movement. By videographing spheroids constituted through mixtures of OVCAR3 cells expressing GFP and RFP, we observed that moruloid spheroids were characterized by a dynamic rearrangement of motile cells within them (Figure 2G; see Video S1). On the other hand, such motility dynamics were not observed in case of blastuloid spheroids (Figure 2H; see Video S2).

We also observed that cells in the periphery of blastuloid (but not moruloid spheroids) showed stretched and flattened nuclei (Figure S4). The absence of motility and flattened appearance of cells on the outer part of spheroids implied that there are external compressive forces that are exerted on spheroids. In fact, examining the surface of ultrastructurally imaged blastuloid spheroids suggested the presence of a coat-like biomaterial that masked the furrows between the cells; such furrows are visible in moruloid spheroids (Fig. 2A and B, v and vi).

The non-fibrillar nature of the coat along with the presence of occasional pores suggested to us that mature spheroids may specifically be covered by a basement membrane (BM)-like coat^21^ on their outer surface (Figure S5). We used immunofluorescence cytochemistry to identify the canonical constituents of BM matrix; non-fibrillar Collagen IV and Laminins (such as Laminin-5) typify the macromolecular organization of BM^21, 22^. We checked for the localization of these proteins in adhesive OVCAR3 monolayers, as well as their moruloid and blastuloid spheroids. Collagen IV localized in the cytoplasm within monolayer OVCAR3 cells but was found to be present specifically in the outer surface of both moruloid and blastuloid spheroids (Figure 3A). Pan-Laminin signals, on the other hand, while also observed in cytoplasm of monolayer and moruloid OVCAR3 cells, localized to the outer surface of blastuloid spheroids (Figure 3B). Similar localization to pan-Laminin was also observed for Laminin-5 (Laminin-332) (Figure S6). The early localization of Collagen IV on the outer surface of moruloid spheroids also suggest that the protein may have a key laminin-independent role in initiating BM formation during spheroidogenesis (this has recently been observed in C. elegans pharyngeal BM formation^23^). Staining with NCAM for blastuloid spheroids confirmed that antibodies could penetrate to the inner cells within such specimen (Figure S7)). We also found Collagen IV and pan-Laminin signals on the outer surface of the patient-derived lumen containing spheroids (Figure 3C and D). Altogether, the formation of a BM-like matrix coat was coincident with the acquisition of blastuloid organization by mature spheroids.

**Figure 3:**
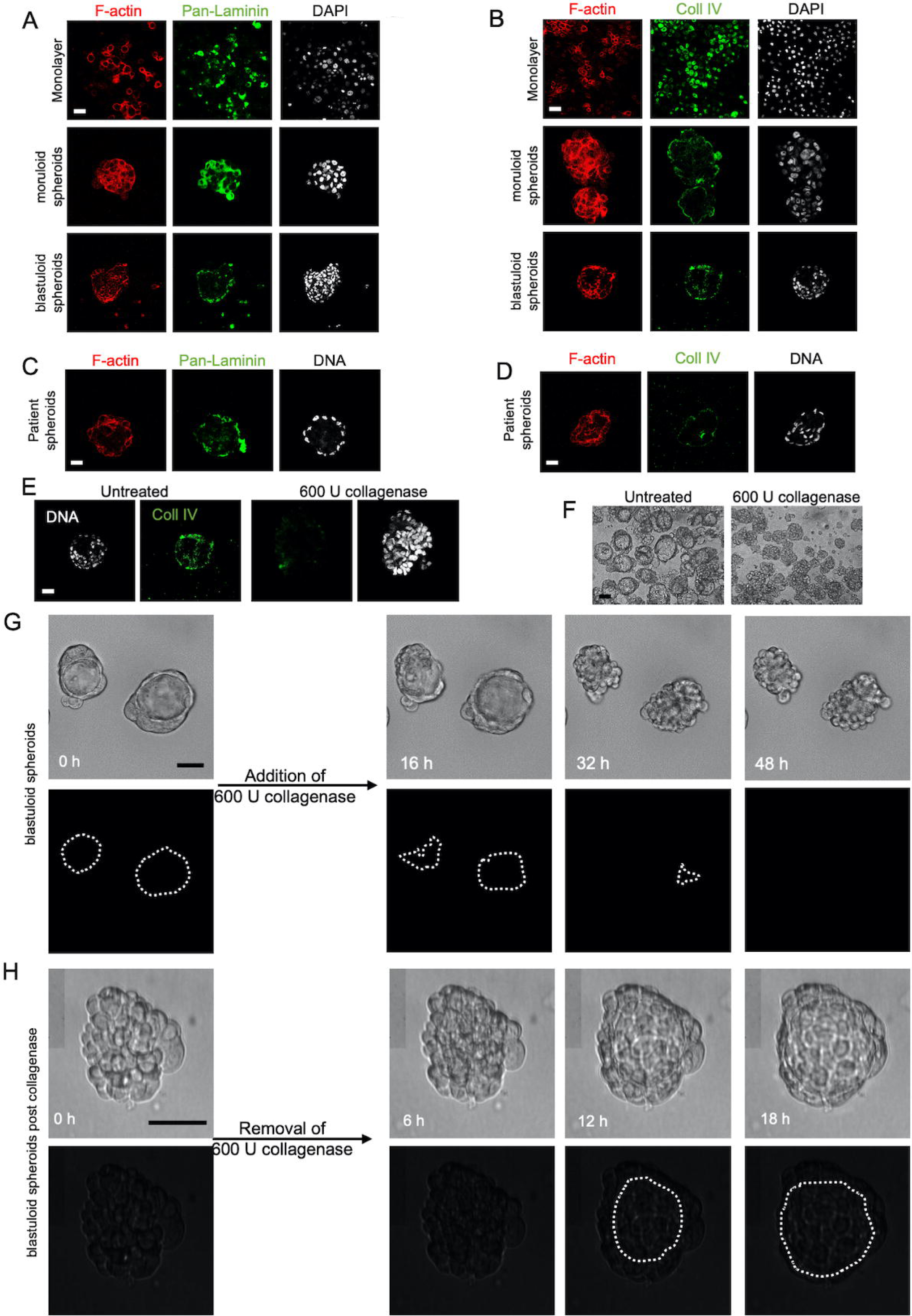
The presence of basement membrane-like ECM coat correlates with lumen formation in spheroids. (A-B) Laser confocal photomicrographs showing Collagen IV (green) (A) and pan-Laminin (green) (B) localization using indirect immunofluorescence in monolayers (top row) moruloid- (middle row) and blastuloid- (bottom row) spheroids from OVCAR3 cells counterstained for F-actin with phalloidin (red) and DNA with DAPI (white). (C-D) Laser confocal photomicrographs showing Collagen IV (green) (C) and pan-Laminin (green) (D) localization using indirect immunofluorescence in patient spheroids counterstained for F-actin with phalloidin (red) and DNA with DAPI (white). (E) Laser confocal photomicrographs stained for Collagen IV (green) and DNA (DAPI; white) in untreated control OVCAR3 spheroids (left) and upon treatment with Collagenase IV (right). (F) Phase contrast photomicrographs showing the morphologies of spheroids with no treatment (control, left) and upon treatment with Collagenase IV (right). G) Bright field photomicrographs taken at 0 hr, 16 hr, 32 hr and 48 hr from time-lapse videography of blastuloid OVCAR3 spheroids initiated after addition of collagenase IV (see Video S3) White dotted lines in the black background highlight the changes in the contour of lumen in G. (H) Bright field photomicrographs taken at 0 hr, 6 hr, 12 hr, 18 hr from time-lapse videography of OVCAR3 spheroids pretreated with Collagenase IV with videography initiated after the removal of Collagenase IV (see Video S4). White dotted lines in the black background highlight the changes in the contour of lumen in H. Scale bar for A-H: 50 μm. Objective 20X.

To investigate, whether the BM coat is involved in imparting stability to the morphology of the mature ovarian cancer spheroid, we treated them with Collagenase IV (in presence and absence of its quencher, FBS). We confirmed the removal of matrix coat upon staining collagenase-treated spheroids for Collagen IV and evincing a decreased signal, when compared with control spheroids (Figure 3E). The removal of matrix coat was additionally confirmed using scanning electron microscopy, where we also noticed a reversal in compaction to a moruloid appearance with a grape-like contour (Figure S8). Such changes in spheroidal morphology were also appreciable in phase contrast microscopy, wherein we detected an accompanying loss of lumen (Figure 3F). We also tracked the process using time-lapse videomicrography, wherein a gradual and progressive depletion of lumen within spheroids (and a reversal of surface compaction) could be observed upon addition of collagenase (Figure 3G; dotted line marks the boundary of the lumen in 3G; Video S3). Upon washing the collagenase away and re-culturing the spheroid in serum-free medium, we noticed a re-transition of the moruloid morphology to a blastuloid phenotype, with re-emergence of lumen and compaction of the surface within 24 hours (Figure 3H; dotted line marks the boundary of the lumen in 3h; Video S4). In fact, intercellular rearrangement and cell motility seen only in moruloid spheroids, also re-emerged in blastuloid spheroids upon debridement of BM by collagenase (Video S4). Our findings suggest that the BM coat mediates the transition of spheroids from an early moruloid- to a blastuloid-phenotype in a highly dynamic and reversible manner.

We next asked if the BM coat also regulates the size and cellular constitution of spheroids. In order to investigate this question, we stably expressed GFP and RFP in separate populations of OVCAR3 cells. In the first experiment, we separately cultured moruloid and blastuloid GFP-expressing spheroids in suspension in the presence of RFP-expressing single OVCAR3 cells. Within 24 hours, we observed RFP cells inside moruloid spheroids but not in blastuloid spheroids (Figure 4A and 4B). In the second experiment, we co-cultured, in suspension, moruloid GFP-expressing and RFP-expressing spheroids. Early spheroids were able to coalesce and give rise to bigger spheroids within 24 hours of coculture (Figure 4C). When this same experiment was performed with mature spheroids, no coalescence was observed; mature spheroids remain sequestered without coalescing (Figure 4D). However, when the above experiment was repeated with blastuloid spheroids that had been treated with Collagenase IV (and their BM coat debrided), there was a partial recapitulation of early spheroidal phenotype: we noticed the presence of RFP cells within BM-less GFP-expressing spheroids (Figure 4E). Coalescence was relatively uncommon although observed in some cases (Figure 4f). Next, we asked whether, in addition to morphogenetic stability, the BM coat of spheroids also regulates their ability to attach to biological (cellular and matrix) substrata.

**Figure 4:**
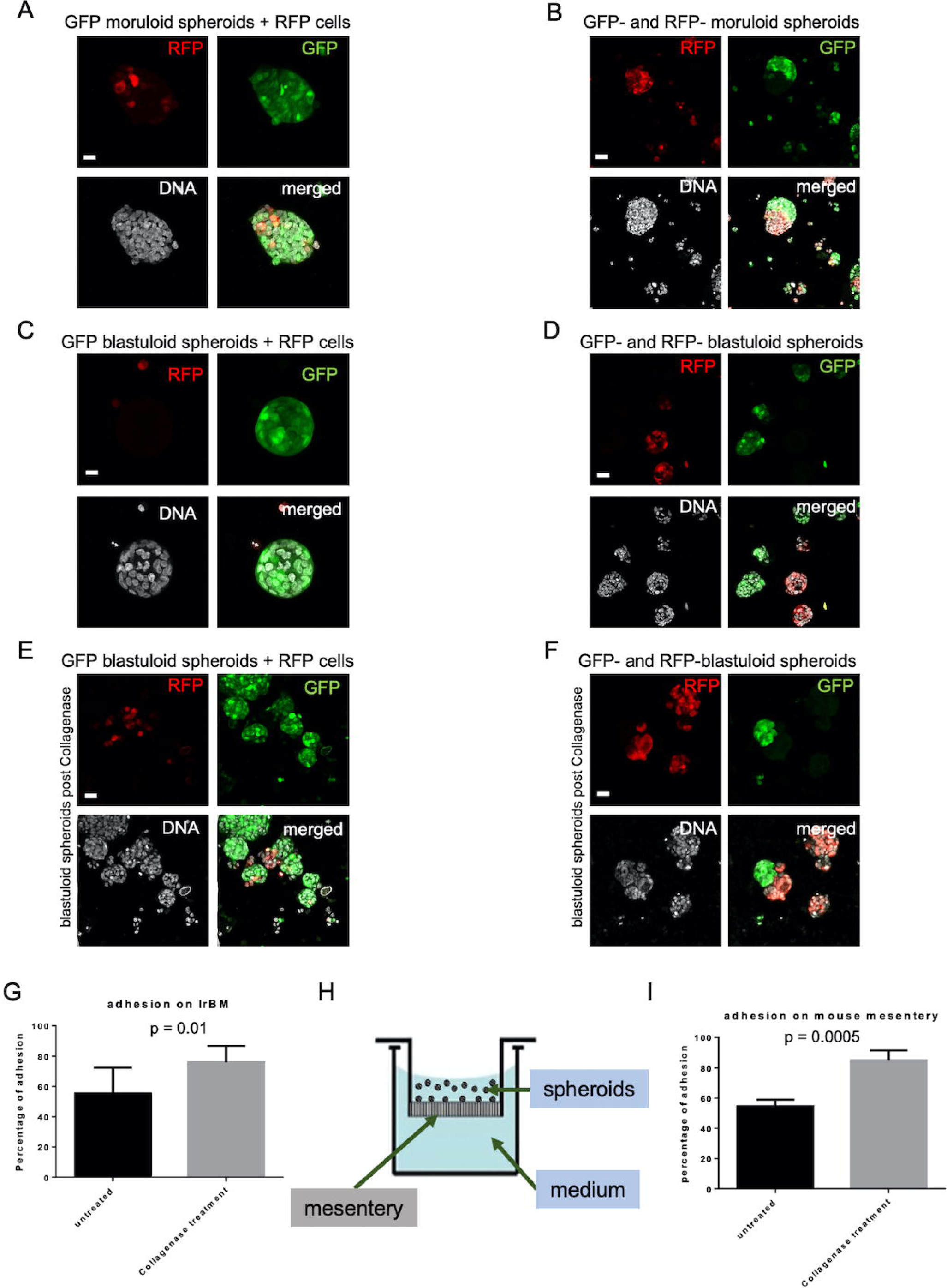
Blastuloid spheroids are morphogenetically more stable than moruloid spheroids. (A-B) Laser confocal photomicrographs of moruloid (A) and blastuloid (B) spheroids expressing GFP, which were cultured with single cells expressing RFP for 24 h and counterstained for DNA (DAPI; white) (C-D)) Laser confocal photomicrographs of spheroids initially formed from separate suspensions of GFP- and RFP-expressing OVCAR3 cells and then cultured together for 24 h and counterstained for DNA (DAPI; white) (E) Laser confocal photomicrographs of blastuloid spheroids expressing GFP, pretreated with Collagenase IV and then cultured with single OVCAR3 cells expressing RFP for 24 h and counterstained for DNA (DAPI; white) (F) Laser confocal photomicrographs of blastuloid spheroids expressing GFP, pretreated with Collagenase IV and then cultured with blastuloid spheroids expressing RFP (also pretreated with Collagenase IV for 24 h and counterstained for DNA (DAPI; white). (G,I) Bar graphs showing relative adhesion of blastuloid spheroids, untreated (left) and pretreated with Collagenase IV (right) when cultured on top of laminin-rich BM scaffolds (G), and 4-6 week BALB/c murine mesenteries that are placed as substrata using transwells (schematic, H) and adhesion studied (I). Bars represent means +/- S.E.M. Scale bar for A-F 50 μm. Objective 20X. Significance was measured using unpaired t-test with Welch’s correction.

We treated spheroids with collagenase IV and cultured them in suspension on top of laminin-rich BM (lrBM)- and Type 1 Collagen ECM (along with untreated controls). We observed that untreated spheroids attached more rapidly to Collagen I than lrBM in consonance with previous demonstrations^6^. Interestingly, BM coat-debrided spheroids showed a significantly higher adhesion to lrBM matrices than their undebrided counterparts (Figure 4G). Higher adhesion was also observed for collagen I scaffolds although the mean percentages of adhesion were not significantly different from untreated controls (Figure S8). In order to recreate the prospective secondary metastatic microenvironment of ovarian cancer *ex vivo*, we dissected out the mesentery of 4-6-week BALB/c mice and sewed it to the bottom of membrane-less transwells (Figure 4H). Upon adding the ovarian cancer spheroids (with and without BM), we assessed their adhesion to the mesothelial lining. Spheroidal BM coat removal resulted in significantly better adhesion even on murine mesenteries when compared with untreated controls (Figure 4I).

Tumors cells are known to secrete ECM that is proteomically and glycochemically distinct from that synthesized by epithelial and connective tissues^24^. In fact, the expression of ECM proteins and glycans by spheroids constituted from thyroid and glioma cancer cell lines has been demonstrated by previous studies^25-27^. We extend these observations to demonstrate the presence of the BM-constituting proteins in metastatic niche of ovarian cancer patients as well as cell lines. Furthermore, we demonstrate important functions mediated by this morphogenetic trait; the first relates to multicellular organization. The appearance of the BM results in loss in cell movement; compaction and stabilization of intercellular relationships and formation of lumen. Cavitation within multicellular structures has been attributed to the apoptosis of centrally located cells in embryonic and postnatal contexts^28, 29^. In contrast, lumen formation may take place without apoptosis, through a separation of apical and basolateral organization within Marine Darby Canine Kidney cells cultured in collagen^30, 31^. Recent research on the formation of blastocoels within murine blastocysts attributes the latter to microfractures in cell-cell contacts followed by fusion of microlumina, similar to Ostwald ripening^32^. The kinetics of lumen appearance and abrogation in ovarian cancer spheroids upon formation and removal of BM respectively suggests an apoptosis-independent mechanism. This is further strengthened by its concurrence with stoppage of intercellular rearrangements that may be detrimental to the establishment of cell-cell contacts necessary for both polarity- and microfracture-dependent mechanisms.

The literature on size regulation in spheroids is scarce. The self-limiting behavior of spheroids has been proposed to depend on proliferation or diffusion of nutrients^33, 34^. Besides proliferation, spheroids may also grow through interspheroidal coalescence^35, 36^. Our observations show that the gradual formation of the BM may act as a barrier to coalescence and incorporation of new cells from the suspended milieu. In doing so, the BM formation also facilitates the spheroids to adopt a dual morphological phenotype with distinct rheological properties: coalescence of BM-coatless moruloid spheroids is prognostic of liquid-like behavior^37^, whereas BM-containing blastuloid spheroids are relatively more solid-like with minimal compositional rearrangement). Such morphological dichotomy might significantly impact how spheroids traverse through peritoneal spaces during metastasis.

The colonization of secondary metastatic sites has long been interpreted through the framework of the ‘seed-and-soil’ hypothesis^38^. Recent research furthers the hypothesis by proposing that premetastatic niche often carries with it agents such as activated fibroblasts (that can act as the soil) that facilitate the colonization^39^. Our results indicate that the ‘soil’ carried may be complex and may not necessarily be inducive: the BM coat of blastuloid spheroids actually impedes colonization relative to the moruloid ones. The consequences of such negative regulation could be linked to the ascitic accumulation of spheroidal population, which may induce collagenase secretion by peritoneal mesothelia^39^. Our findings therefore suggest a link between the buildup of the ovarian cancer metastatic niche and its mesenteric colonization, rendering it an important target for future studies.

## Materials and Methods

### Cell culture

The human ovarian cancer cell lines: OVCAR3, OVCAR4, SKOV3 were a kind gift from Professor Rajan R. Dighe, IISc. OVCAR3 and OVCAR4 were maintained in Dulbecco’s modified Eagle’s medium (DMEM) (HiMedia AL007A) supplemented with 10-20 % fetal bovine serum (FBS) (Gibco, 10270) and antibiotics in a humidified atmosphere of 95% air and 5% CO2 at 37°C. The SKOV3 cell line was maintained in McCoy’s 5A medium (HiMedia AL275A) supplemented with 10% FBS and antibiotics. Another ovarian cancer cell line G1M2 was a kind gift from Dr. Sharmila A. Bapat, NCCS. G1M2 cell line was maintained in RPMI (Roswell Park Memorial Institute) (HiMedia AL162A) supplemented with 10% FBS and antibiotics.

### Spheroid culture

Spheroids were cultured on dishes coated with 3% poly-2-hydroxyethyl methacrylate (polyHEMA) (Sigma, P3932), which was dissolved in 95% absolute ethanol overnight in magnetic stirrer at room temperature (RT). PolyHEMA coating prevented cell attachment and allowed for spheroid formation in suspension. Cells were seeded according to the requirement for experiment in defined medium: DMEM:F12 (1:1) supplemented 0.5 μg/ml hydrocortisone (Sigma, H0888), 10 μg/ml insulin (Sigma, I6634), and 10 ng/ml recombinant human epidermal growth factor (hEGF) (HiMedia, TC228). Spheroids formed in the suspension were visualized using phase contrast or brightfield microscopy. Spheroids were collected from the cultures by centrifugation at 800-1200 rpm for 5 minutes.

### Clinical samples

Ascites obtained from the paracentesis of patients with ovarian cancer was provided by Sri Sankara Cancer Hospital with due ethical clearance. Patient spheroids were cultured on tissue culture-treated polystyrene substrata/ polyHEMA coated dish using Dulbecco’s modified Eagle’s medium (DMEM) (HiMedia AL007A) - supplemented with 10-20 % fetal bovine serum (FBS) (Gibco, 10270) and antibiotics - and the ascites fluid of the patient in a 1:1 ratio, and subsequently in DMEM/10% FBS alone. Spheroids were then collected from the cultures by centrifugation at 800-1200 rpm for 5 minutes.

### Immunostaining and image acquisition

Cells were subjected to fixation using 3.7% formaldehyde (Fisher Scientific, 24005) at 4°C for 20 min. Following this, the cells were taken for further processing or stored in 1X Phosphate buffered saline (PBS) at 4°C. Spheroids were dried and fixed on 8 well chambered cover glass by placing them on dry bath at 37 °C. After fixation, permeabilization was achieved using 0.5% TritonX-100 (HiMedia, MB031) for 2 hours in monolayer cultures and spheroids. Effective permeabilization is needed for entry and uniform exposure of the antibodies. Blocking was achieved by treating with blocking solution (PBS with 0.1% TritonX-100 and Bovine serum albumin (BSA) (HiMedia, MB083) for 45 mins at RT. Primary antibody incubation was carried out overnight at 4°C. This was followed by washes using 0.1% TritonX-100 in PBS (5 minutes X 3). Secondary antibody incubation was done at RT for 2 hours under dark conditions. DAPI (Thermo Fischer Scientific, D1306) was added to the samples and washed away after 15 minutes. Subsequent processing was carried in the dark. This was followed by washes using 0.1% TritonX-100 in PBS (5 minutes x 3). Images were captured in 20X using Carl Zeiss LSM880. Images were processed and analyzed using ZENLITE software.

### Removal of spheroidal matrix using collagenase

Spheroids were cultured in 35 mm dish by seeding 1.5 ×10^5^ cells for 1 week. Collagenolysis of mature spheroids was performed using 600 U of collagenase IV (Sigma, C5138) with or without 20% FBS for 24 hours. FBS was used as a quenching control for collagenolytic activity. Both control and collagenolysed spheroids were fixed and processed for immunocytochemistry (ICC) or scanning electron microscopy (SEM). Later, to check the effect of collagenolysis on spheroidogenesis, spheroids were cultured in the presence of collagenase and imaged in time-lapse using a brightfield epifluorescence microscope (Olympus IX73).

### Scanning electron microscopy

Monolayer cells and spheroids were fixed using 2.5% glutaraldehyde (Amresco, 0875) overnight followed by three PBS (5 minutes each) washes to remove excess fixative. This was followed by five washes with water to remove salts. Dehydration was followed using different grades of ethanol (30%, 50%, 70%, 90%, 100%). Spheroids were then put on 1N HCl-treated coverslips and allowed to dry completely at room temperature. Cellular monolayers were directly seeded on top of the coverslip. Imaging was done using ESEM Quanta.

### Adhesion assay on murine mesenteries

BALB/c female mice (4-6 weeks old) were used for adhesion experiments. Mice were sacrificed by cervical dislocation and upon surgically dissecting the abdomen, their mesentery were strung to the lower ends of transwell inserts (Boyden chambers without the membranes) using surgical thread. The transwells containing mesentery were placed in a 24-well tissue culture plate and defined medium added both into the transwell insert and below the insert into wells of the plate. Spheroids were then added onto the upper layer of the mesentery on the insert; making sure the media does not spill from the insert into the well and adhesion studied through brightfield microscopy after washing at fixed intervals.

### ECM coating for adhesion assay

8 well chambered cover glasses (Eppendorf, 0030742036) were coated with 50 μg/mL Growth factor reduced basement membrane matrix (Matrigel®) (Corning, 354230) or 1mg/mL rat tail collagen neutralized on ice in the presence of 10X DMEM with 0.1 N NaOH such that the final concentration of the collagen is 1 mg/ml.

### Adhesion assay

Spheroids were cultured for 1 week in polyHEMA-coated 35 mm dishes using defined medium. After 1 week of culture, spheroids were treated with Type IV collagenase (Gibco 17104019) for 24 hours. Collagenase activity was quenched using 10% FBS, followed by PBS washes; untreated spheroids were used as control. Spheroids were resuspended in 1 ml defined medium for the assay. 10-20 μl of both untreated and treated spheroids suspension were put on top of ECM-coated chamber wells and isolated mesenteries in transwells. The number of spheroids seeded was counted, allowed to attach for 6 hrs by incubating them inside humidified 37 °C 5% CO_2_ incubator. After 6 hours of incubation, the medium was replaced with fresh PBS to wash away unattached spheroids and the adhered ones were counted. The percentage of adhesion was calculated by dividing the amount of spheroids attached to the substrata by the total number of spheroids seeded.

### Bright field time lapse microscopy

Spheroids from cell lines were cultured for 24 hours and 1 week by seeding 1.5 x 10^5^ cells in a 35 mm cell culture dish. At particular time points, spheroids were harvested by centrifugation at 1200 rpm for 5 minutes, resuspended and put on a drop of 4% noble agar (Sigma-Aldrich, A5431) that was smeared on a glass-bottomed chamber well, which after some time was flooded with defined medium. Time-lapse imaging was subsequently performed for 48 hrs with 15 minutes interval using a Tokai Hit stage-top incubator with image acquisition through a Orca Flash LT plus camera (Hamamatsu) on an Olympus IX73 microscope.

### Confocal time lapse microscopy

We established GFP- and RFP-expressing OVCAR3 cell lines using lentiviral transduction. Using these cell lines, we cultured spheroids for 24 hours or 1 week on PolyHEMA-coated 35 mm dishes either individually or in combination -as described in the results section. Spheroids (made from mixtures of RFP- and GFP-expressing cells) were then harvested and immobilized for time lapse microscopy on a bed of 4% noble agar following similar protocol described above. Time lapse imaging was performed using LEICA SP8 confocal microscope for 6 hours with 15 minutes interval. Data was analyzed using LASX Leica software.

## Acknowledgments

This work was supported by the Wellcome Trust/DBT India Alliance Fellowship Grant [IA/I/17/2/503312] awarded to RB. RB would additionally acknowledge support from the DBT IISc partnership program (BT/PR27952/INF/22/212/2018) and the Institute of eminence grant (IE/CARE-19-0319). JL acknowledges IISc for fellowship. The studies described here were carried out according to the guidelines of the institutional review board and in agreement with the ethical guidelines of Sri Shankara Cancer Hospital and Research Centre, and the Indian Institute of Science (IISc) after obtaining informed consent from the patients. We would also like to thank the Biological Division Imaging facility and the Advanced Facility for microscopy and analysis (AFMM) for help with microscopy

## Figure legends

**Video S1: Moruloid OVCAR3 spheroid showing intercellular movement**

**Video S2: Blastuloid OVCAR3 spheroid showing no intercellular movement**

**Video S3: Progressive loss of lumen and compaction in blastuloid spheroids upon addition of Collagenase IV**

**Video S4: Restoration in lumen and compaction in Collagenase IV treated blastuloid spheroids upon washing away enzyme**.

---

**Figure S1:**
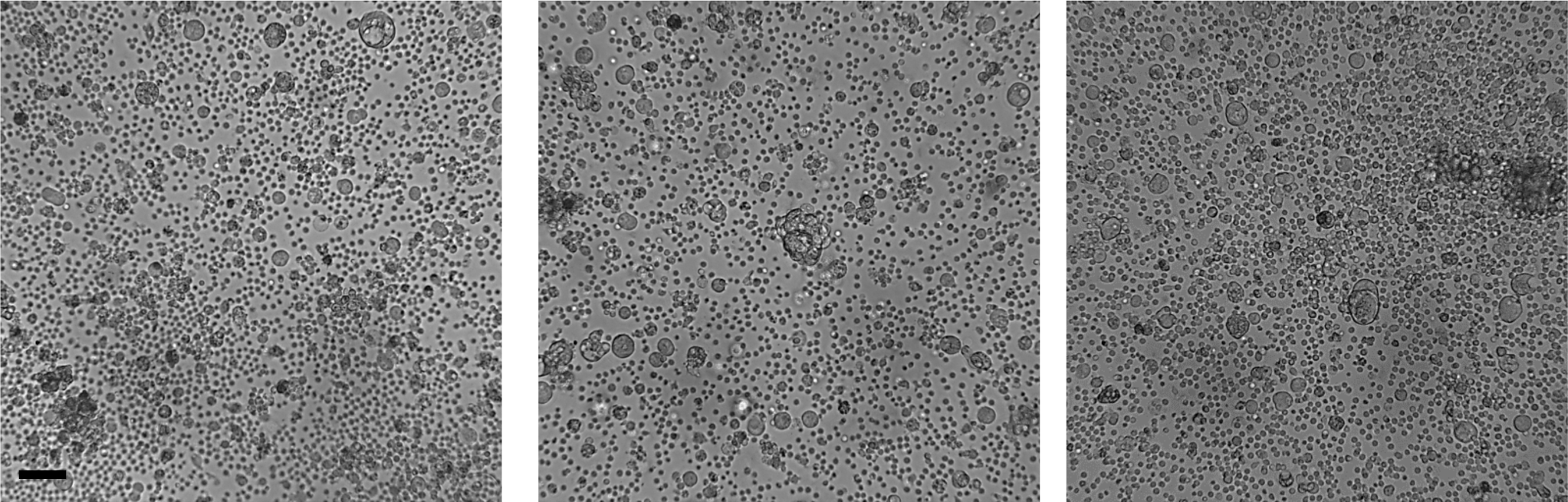
Bright field images of patient spheroids showing lumen-less, lumen containing spheroids and single cells in a same sample. Scale bar: 50 μm. Objective 20X.

**Figure S2:**
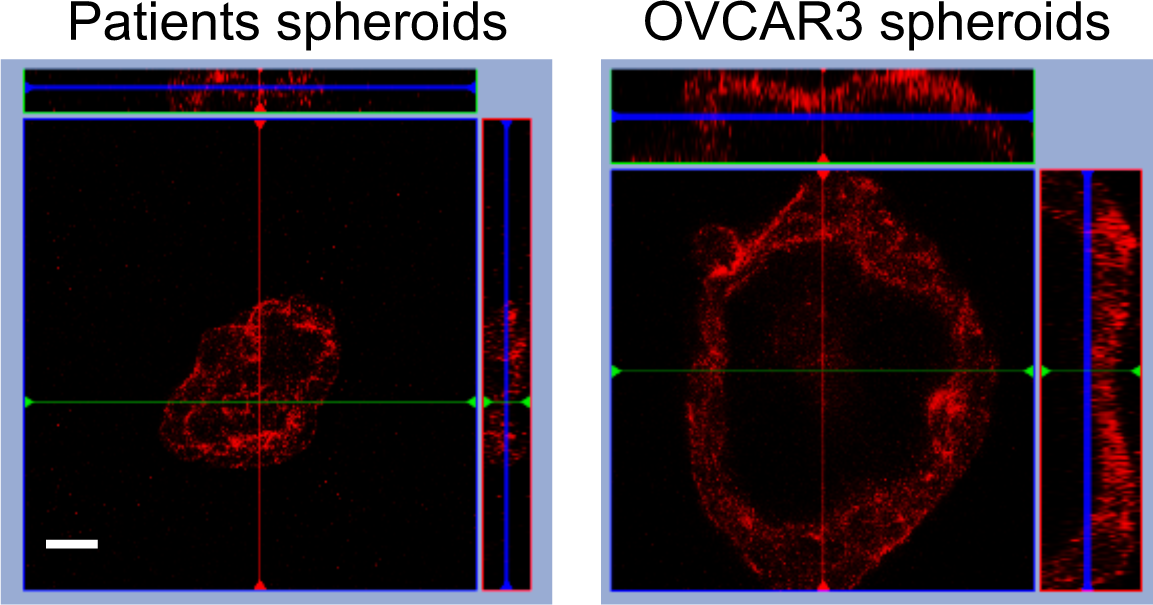
Laser confocal photomicrograph middle stack-orthogonal axes confirming the lumen present in the spheroids of patients (left) and OVCAR3 (right). Scale bar: 50 μm. Objective 20X.

**Figure S3:**
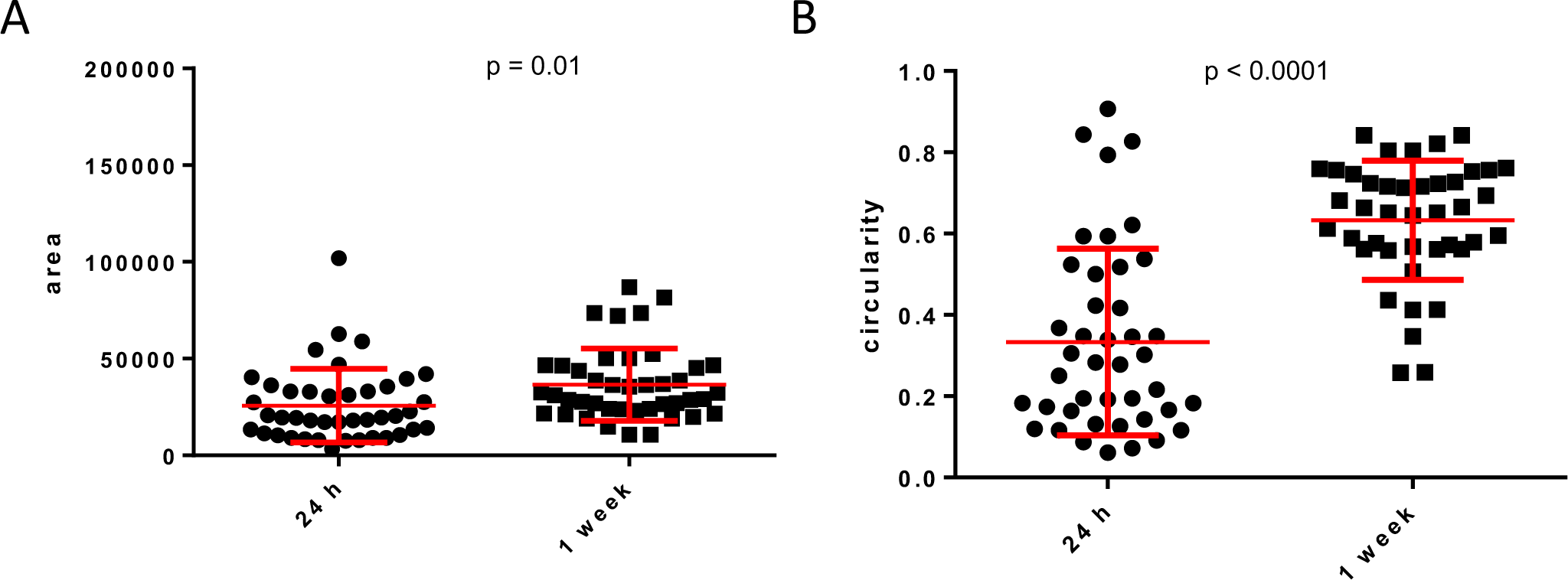
Quantification of G1M2 moruloid and blastuloid spheroids, showing the increased size (A) and increased in circularity (B) during spheroidal maturation.

**Figure S4:**
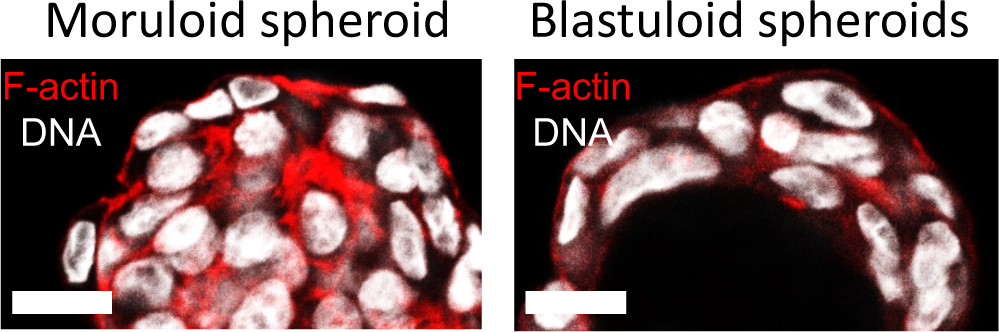
Laser confocal photomicrograph zoom-in images of OVCAR3 moruloid (left) and blastuloid (right) spheroids showing the transition of nuclear shape during spheroids maturation. Nuclear size has flattened in blastuloid spheroids. F-actin (red), nucleus (white). Scale bar: 50 μm. Objective 20X.

**Figure S5:**
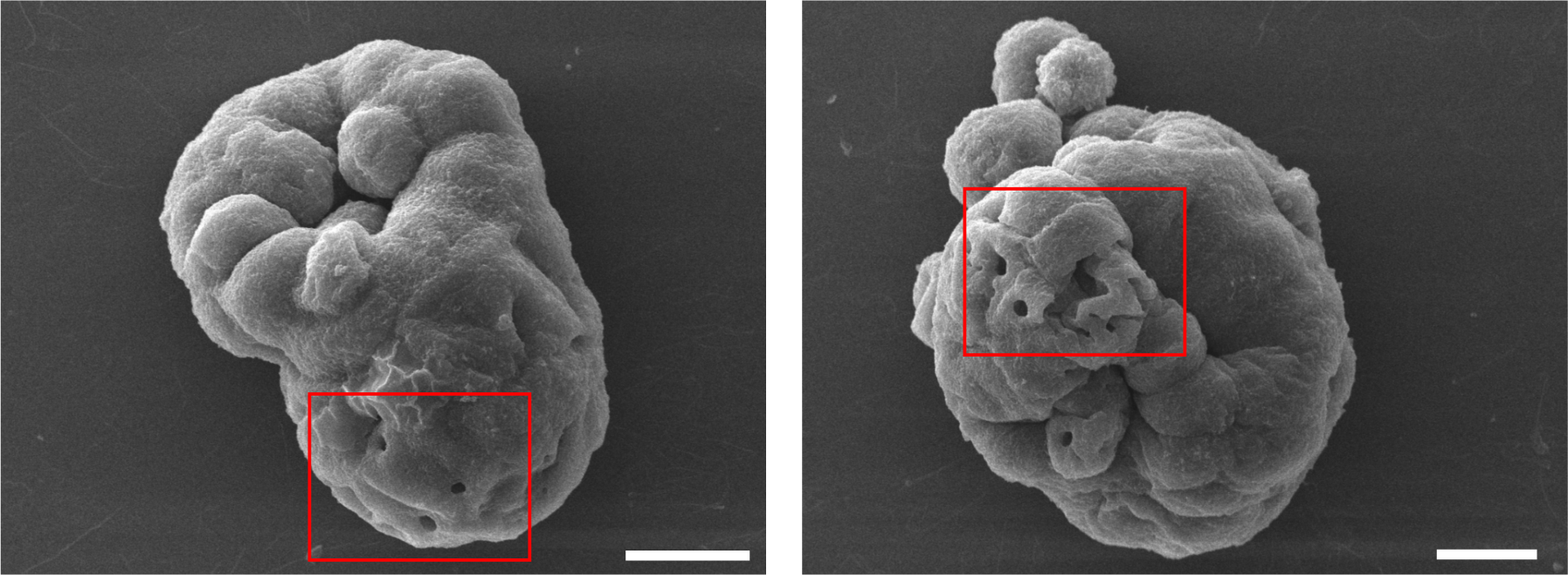
SEM photomicrographs of G1M2 spheroids showing porous basement membrane like matrix on the surface (highlighted in red box). Scale bars = 50 μm.

**Figure S6:**
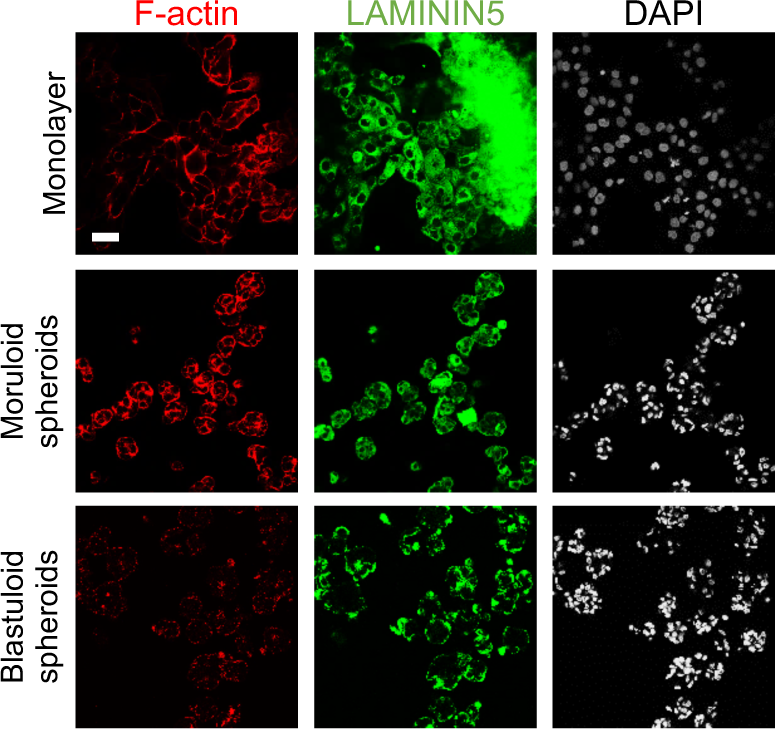
Laser confocal photomicrographs showing Laminin5 (green) localization using indirect immunofluorescence in monolayers (top row) moruloid-(middle row) and blastuloid-(bottom row) spheroids from OVCAR3 cells counterstained for F-actin with phalloidin (red) and DNA with DAPI (white). Scale bar: 50 μm. Objective 20X.

**Figure S7:**
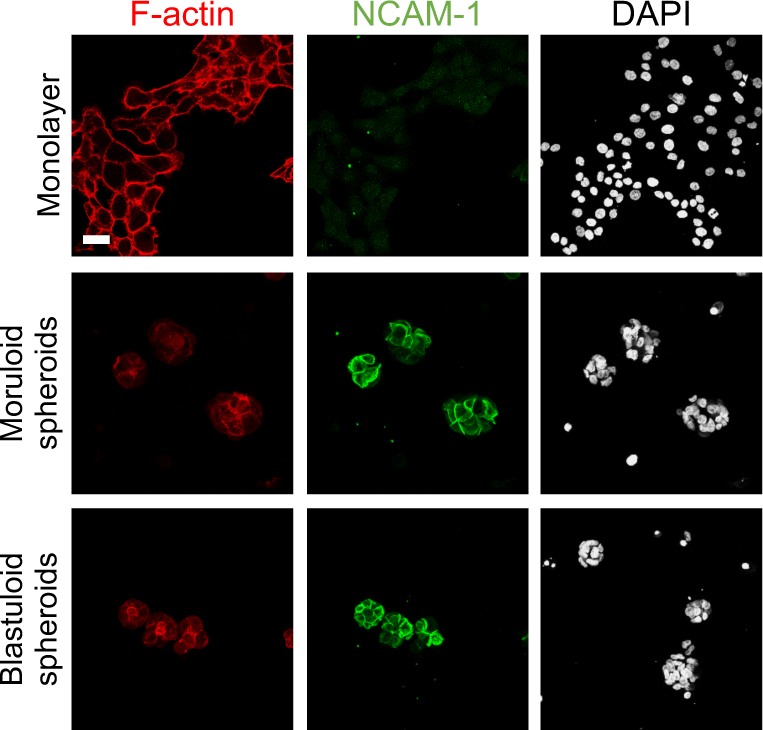
Laser confocal photomicrographs showing NCAM1 (green) localization using indirect immunofluorescence in monolayers (top row) moruloid-(middle row) and blastuloid-(bottom row) spheroids from OVCAR3 cells counterstained for F-actin with phalloidin (red) and DNA with DAPI (white). Scale bar: 50 μm. Objective 20X.

**Figure S8:**
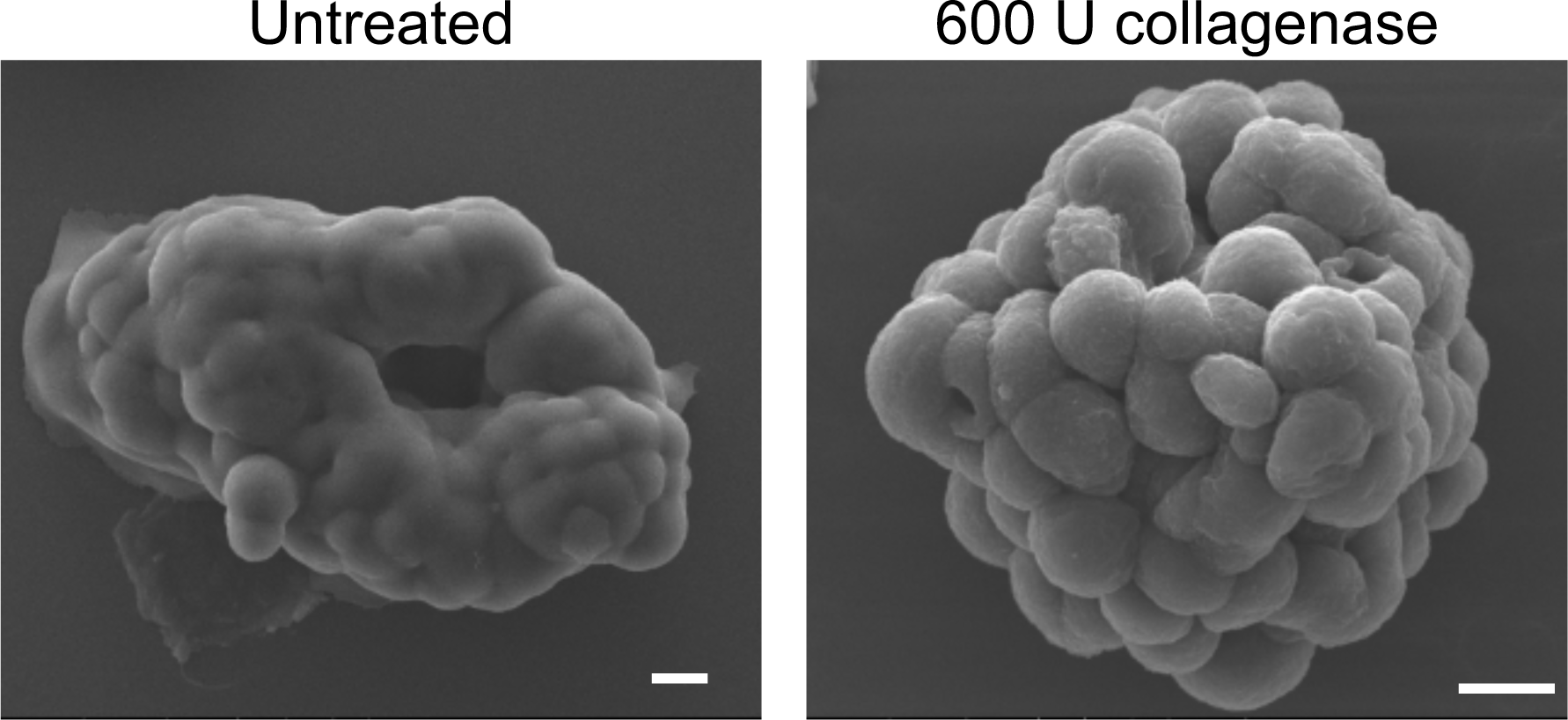
SEM photomicrographs of OVCAR3 spheroids untreated (left) and 600 U collagenase treated (right), confirming removal of matrix. Scale bar = 10 μm

**Figure S9:**
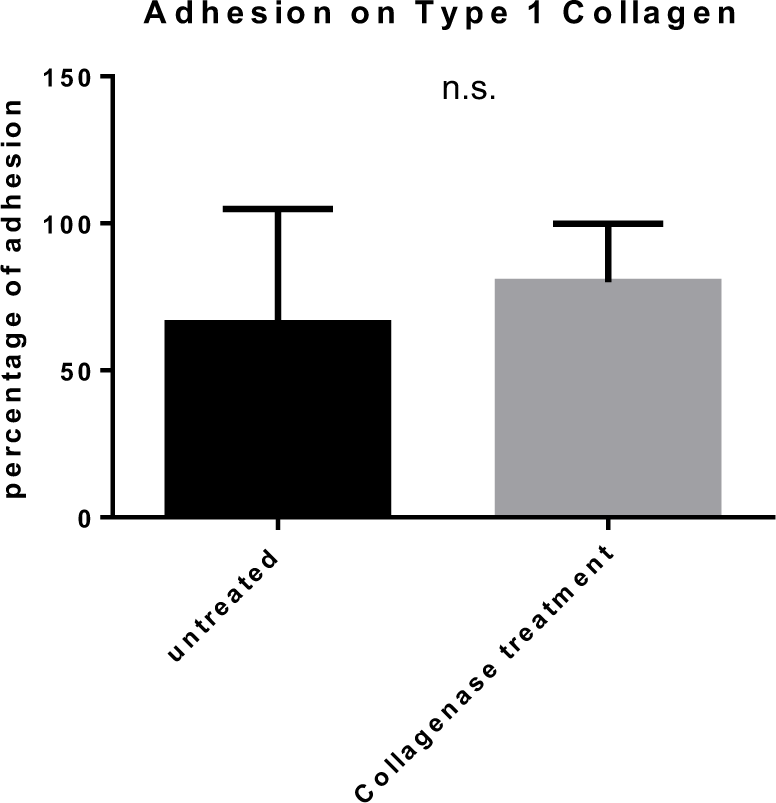
Bar graph showing relative adhesion of blastuloid spheroids, untreated (left) and pretreated with Collagenase IV (right) when cultured on top of Collagen I scaffold. Bars represent means +/- S.E.M. Significance was measured using unpaired t-test with Welch’s correction.

